## Supplementary Material for "Rapid evolution in necromass use under resource limitation reduces persistence in producer-decomposer microbial biospheres"

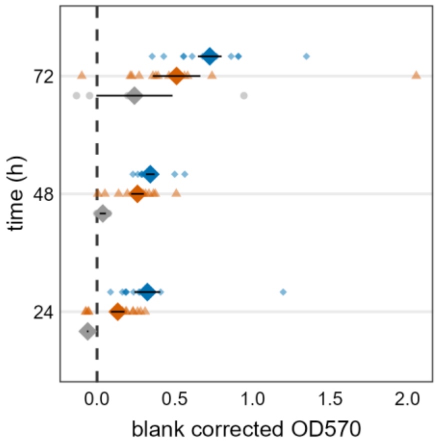


**Supplementary Figure 1.** Biofilm formation capacity at three different cultivation time points (24, 48, 72 hours). Data are derived from blank-corrected OD570 values obtained via the quantitative tissue culture plate (TCP) assay, facilitating comparisons across time points for each treatment group. Color key: gray = WT, orange = HTG, blue = HMG.

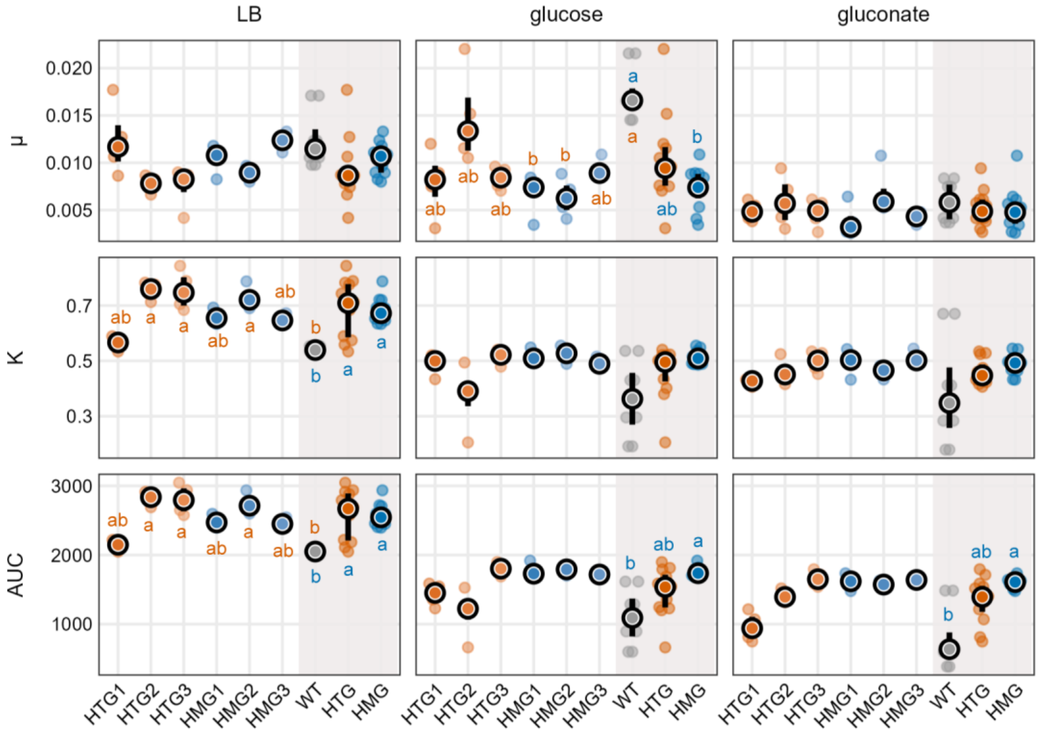


**Supplementary Figure 2.** Comparison of three growth parameters—maximum growth rate (μ), carrying capacity (K), and area under the curve (AUC)—for growth curves of evolved *Escherichia coli* (*E. coli*) MC4100 populations and the wild-type progenitor (WT) on three carbon sources: LB, glucose, and gluconate. Individual data points represent technical replicates; black dots and vertical lines indicate medians and interquartile ranges, respectively. The shaded area on the right illustrates the comparisons between the WT and the respective treatment group to which each population belongs. Orange hues denote populations subjected to monoculture in the HTG (spatially heterogeneous) system, while blue hues denote those from the HMG (spatially homogeneous) system. Note that in this experiment, WT refers to both the wild-type progenitor population and the untreated control group. Orange letters indicate statistically significant differences between individual populations, and blue letters indicate significant differences between treatment groups. Means without a common letter differ significantly (Dunn's test; $\alpha = 5\%$).

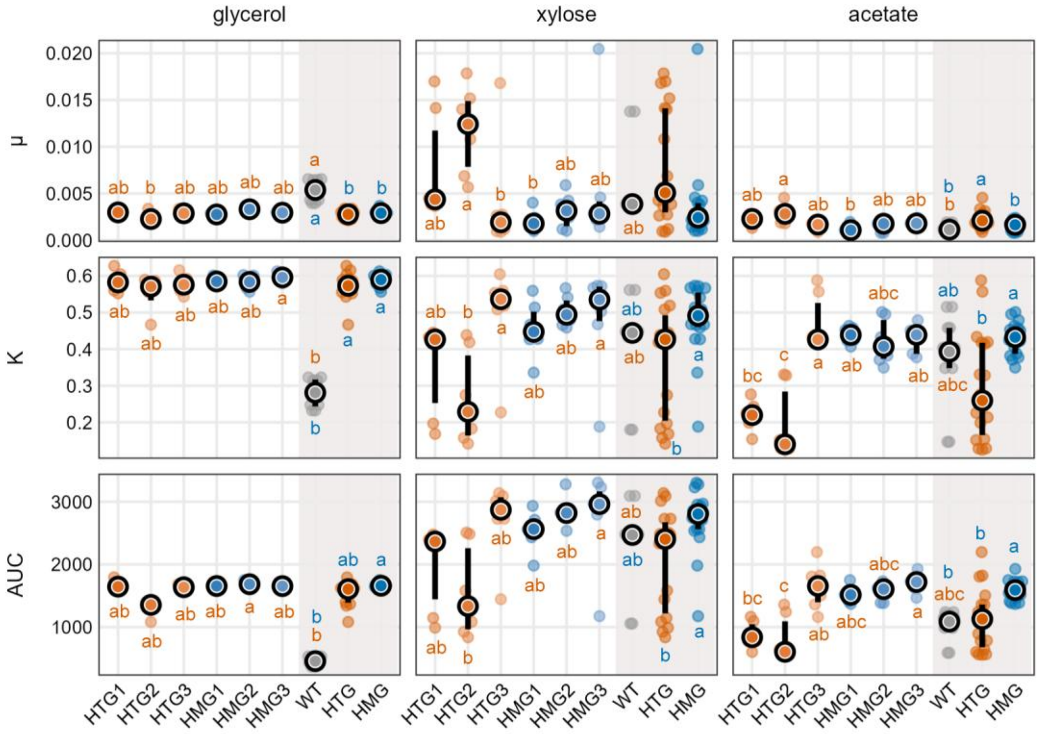


**Supplementary Figure 3.** Comparison of three growth parameters—maximum growth rate (μ), carrying capacity (K), and area under the curve (AUC)—for growth curves of evolved *Escherichia coli* (*E. coli*) MC4100 populations and the wild-type progenitor (WT) on three carbon sources: glycerol, xylose, and acetate. Individual data points represent technical replicates; black dots and vertical lines indicate medians and interquartile ranges, respectively. The shaded area on the right illustrates the comparisons between the WT and the respective treatment group to which each population belongs. Orange hues denote populations subjected to monoculture in the HTG (spatially heterogeneous) system, while blue hues denote those from the HMG (spatially homogeneous) system. Note that in this experiment, WT refers to both the wild-type progenitor population and the untreated control group. Orange letters indicate statistically significant differences between individual populations, and blue letters indicate significant differences between treatment groups. Means without a common letter differ significantly (Dunn's test; $\alpha = 5\%$).

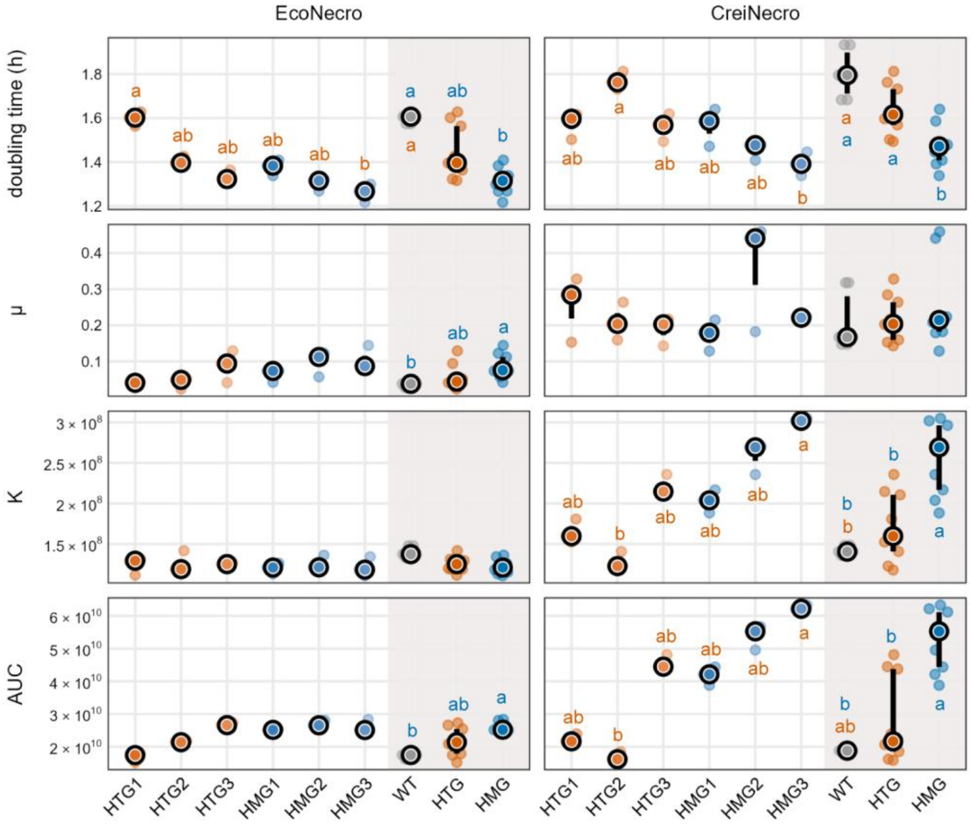


**Supplementary Figure 4.** Comparison of four growth parameters—doubling time, maximum growth rate (μ), carrying capacity (K), and area under the curve (AUC)—for growth curves of evolved *Escherichia coli* (*E. coli*) MC4100 populations and the wild-type progenitor (WT) on EcoNecro and CreiNecro. EcoNecro and CreiNecro denote necromass derived from *E. coli* and *Chlamydomonas reinhardtii*, respectively. Individual data points represent technical replicates; black dots and vertical lines indicate medians and interquartile ranges, respectively. The shaded area on the right illustrates the comparison between the WT and the respective treatment group to which each population belongs. Orange hues denote populations subjected to monoculture in the HTG (spatially heterogeneous) system, while blue hues denote those from the HMG (spatially homogeneous) system. Note that in this experiment, WT refers to both the wild-type progenitor population and the untreated control group. Orange letters indicate statistically significant differences between individual populations, and blue letters indicate significant differences between treatment groups. Means without a common letter differ significantly (Dunn's test; $\alpha = 5\%$).

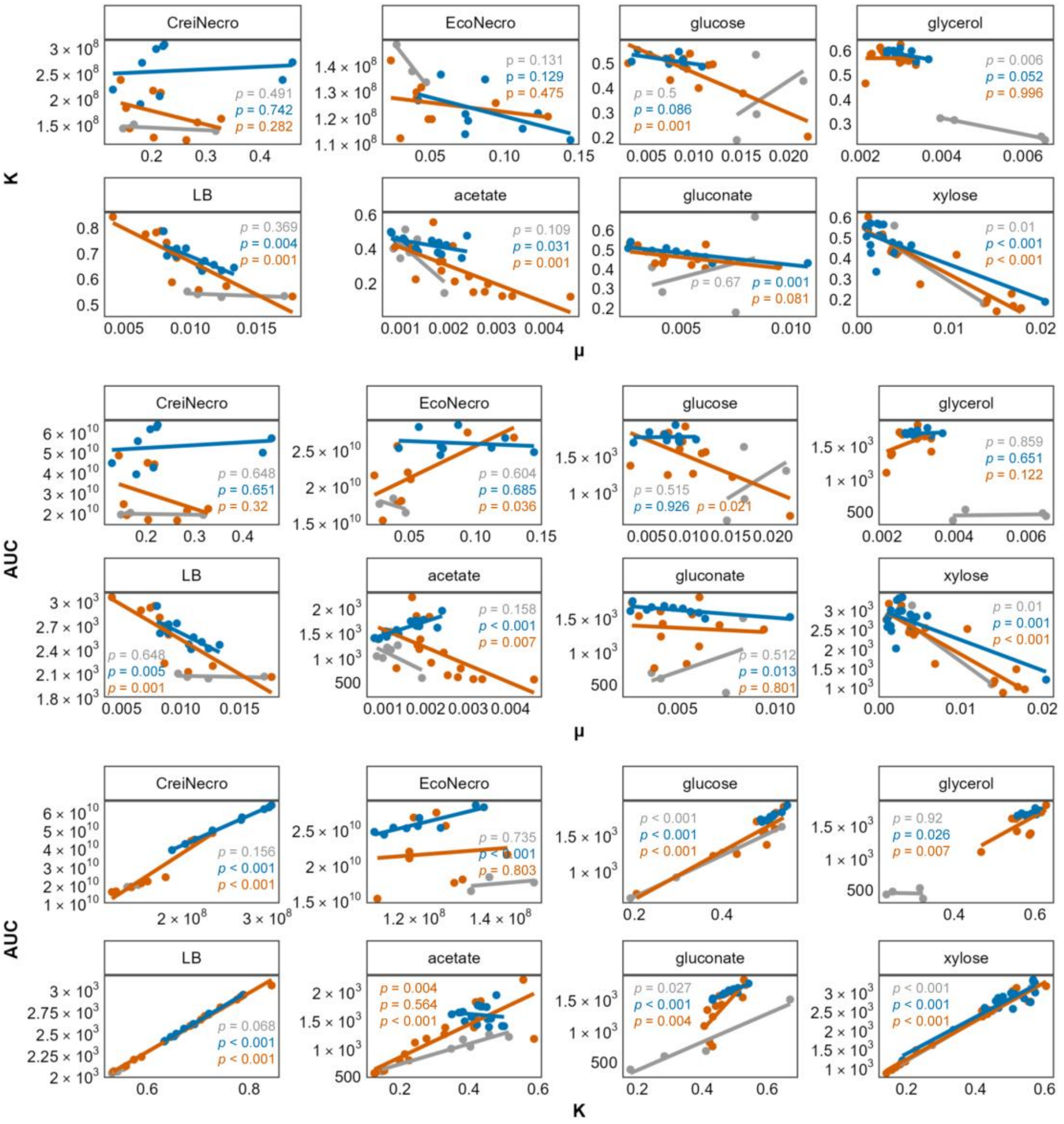


**Supplementary Figure 5.** Pairwise relationships among growth parameters for *Escherichia coli* populations from the three treatments after monoculture, assessed across eight carbon sources. The three treatments are: gray for the untreated wild-type progenitor, orange for populations from HTG (spatially heterogeneous) monoculture, and blue for those from HMG (spatially homogeneous) monoculture. Scatter plots illustrate the distribution between maximum growth rate (μ), carrying capacity (K), and area under the curve (AUC) under different carbon source conditions. Correlations were evaluated and fitted using ordinary least squares (OLS) regression. Top two rows: K vs. μ; Middle two rows: AUC vs. μ; Bottom two rows: AUC vs. K.

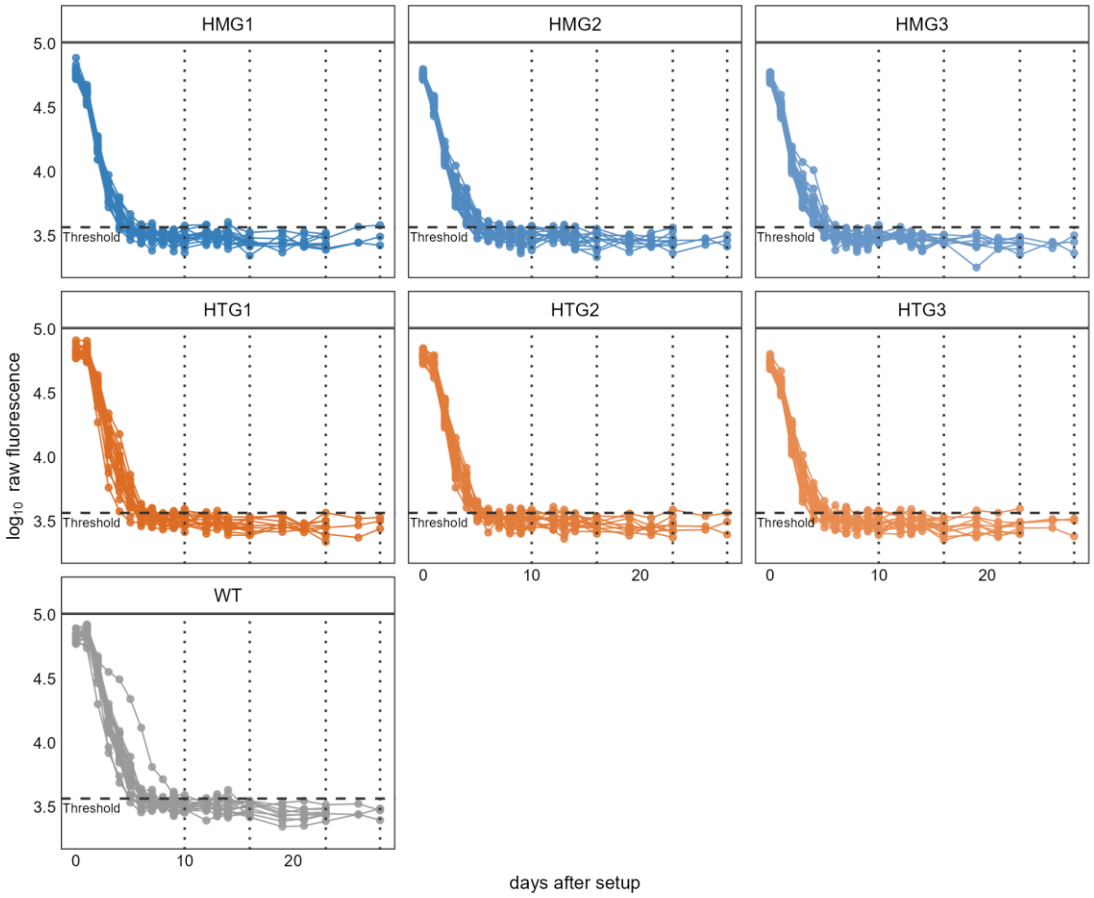


**Supplementary Figure 6.** Non-invasive fluorescence monitoring of co-cultured microbial biospheres over a 28-day period. The horizontal dashed line indicates the detection limit, calculated as the mean fluorescence of the negative control plus three times its standard deviation (SD). Vertical dotted lines mark the four destructive sampling time points at Days 10, 16, 23, and 28. Raw fluorescence measurements were $\log_{10}$-transformed.

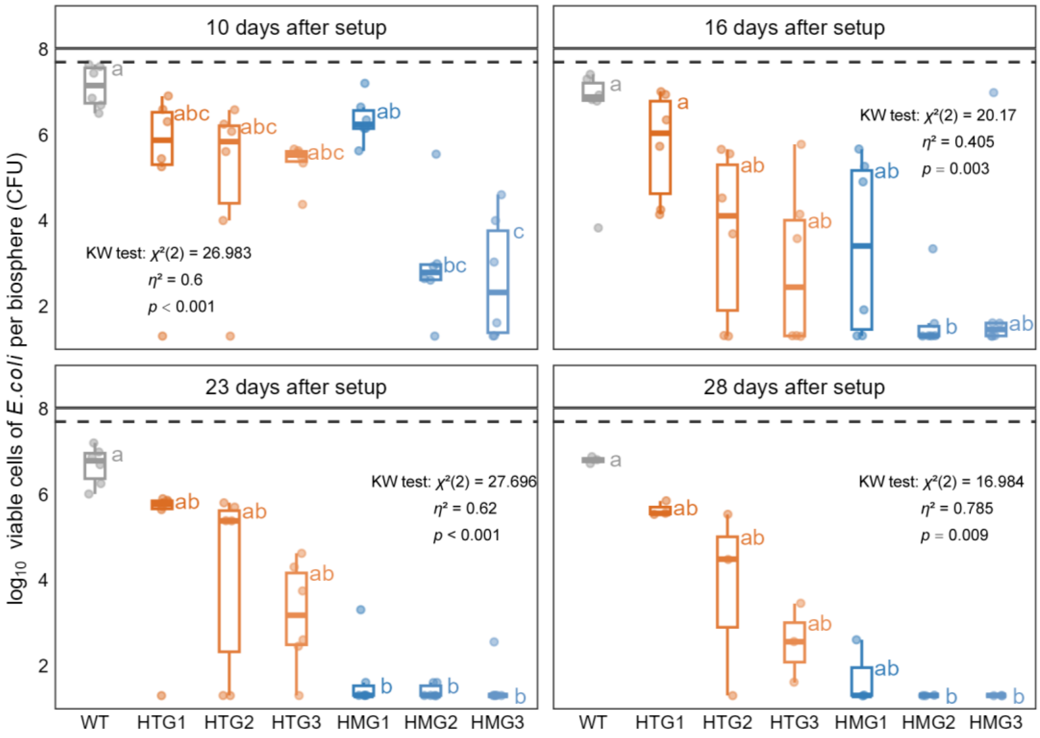


**Supplementary Figure 7.** $Log_{10}$-transformed *Escherichia coli* (*E. coli*) population densities (cells per biosphere) for different *E. coli* populations at the four destructive harvest time points. Gray represents the untreated wild-type progenitor (WT); orange hues denote populations from HTG (spatially heterogeneous) monoculture; blue hues denote populations from HMG (spatially homogeneous) monoculture. At each time point, means not sharing a common letter differ significantly among treatment groups (Dunn's test). The dashed line at the top indicates the initial number of *E. coli* cells inoculated per biosphere.

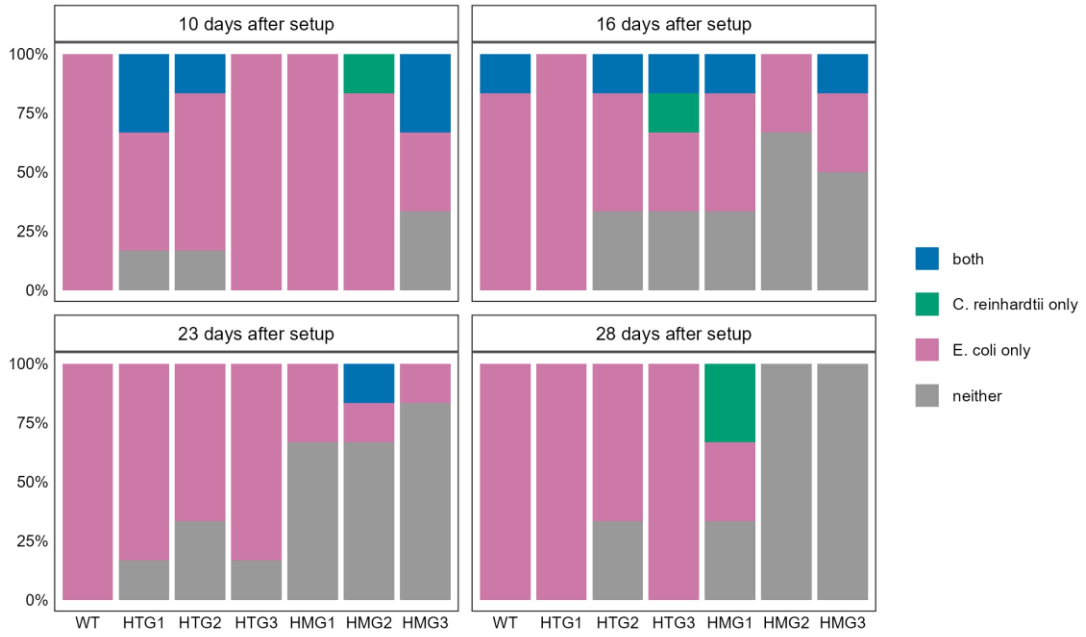


**Supplementary Figure 8.** Percentages of biospheres in each state relative to the total number harvested per time point across the four harvests, for the different *Escherichia coli* (*E. coli*) populations. “both” indicates biospheres where both *E. coli* and *Chlamydomonas reinhardtii* (*C. reinhardtii*) survived; “neither” indicates the opposite. “C. reinhardtii only” and “E. coli only” indicate biospheres where only the alga or the bacterium survived, respectively. WT denotes the untreated wild-type progenitor; the prefixes HTG and HMG denote populations subjected to monoculture in the HTG (spatially heterogeneous) and HMG (spatially homogeneous) systems, respectively.

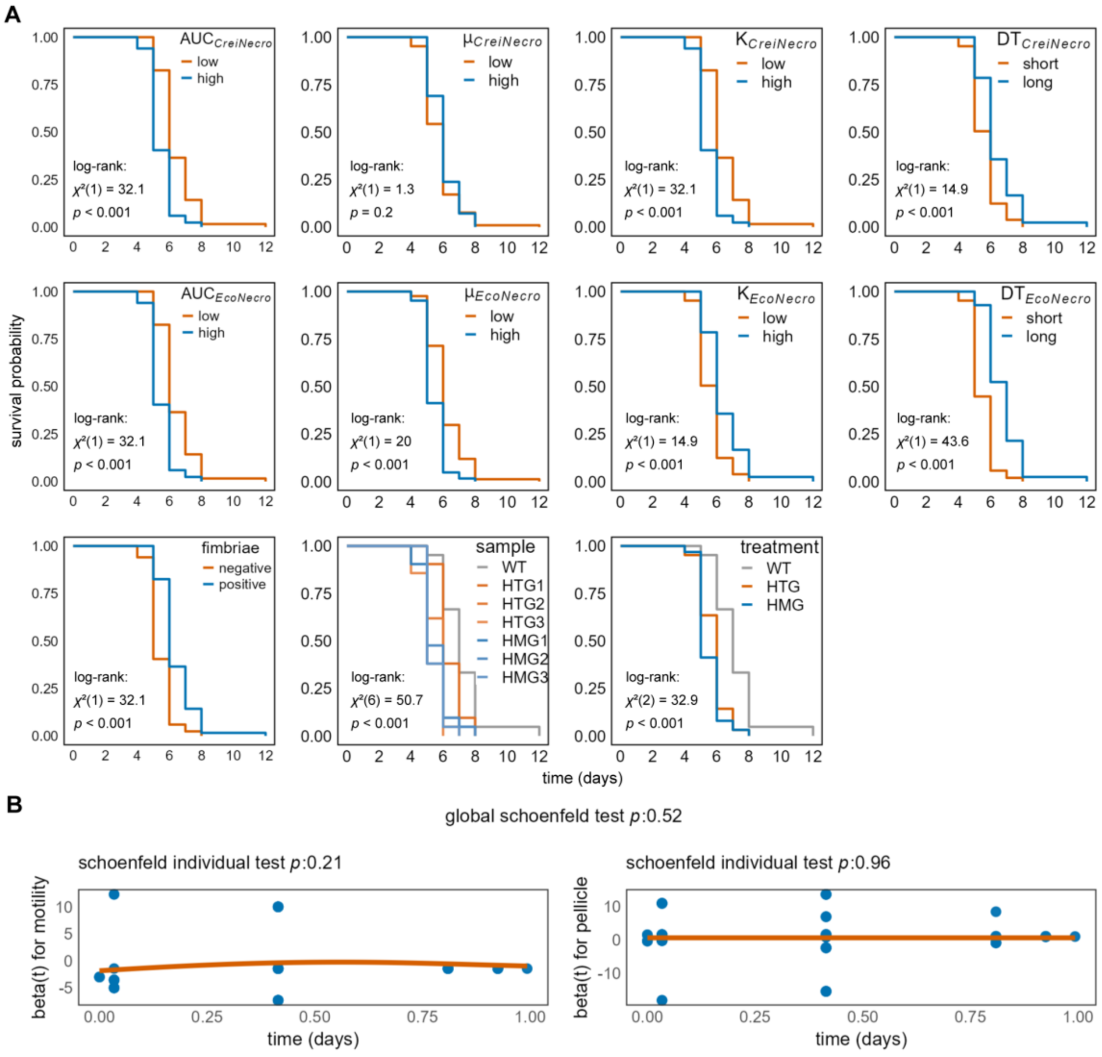


**Supplementary Figure 9. Supplementary results from survival analysis.** (A) Kaplan-Meier survival probability curves for microbial biospheres, stratified by individual sample, treatment group, fimbriae expression, and the growth parameters of growth curves on two types of necromass. Curli fimbriae expression was assessed qualitatively using Congo Red indicator (CRI) agar, with results on a binary scale.[1] CreiNecro and EcoNecro denote necromass derived from *Chlamydomonas reinhardtii* and *Escherichia coli*, respectively. The growth parameters are area under the curve (AUC), maximum growth rate (μ), carrying capacity (K), and doubling time (DT). (B) Schoenfeld residuals versus time, used to test the proportional hazards assumption of the Cox proportional hazards model; a *p*-value > 0.05 indicates that the assumption holds.

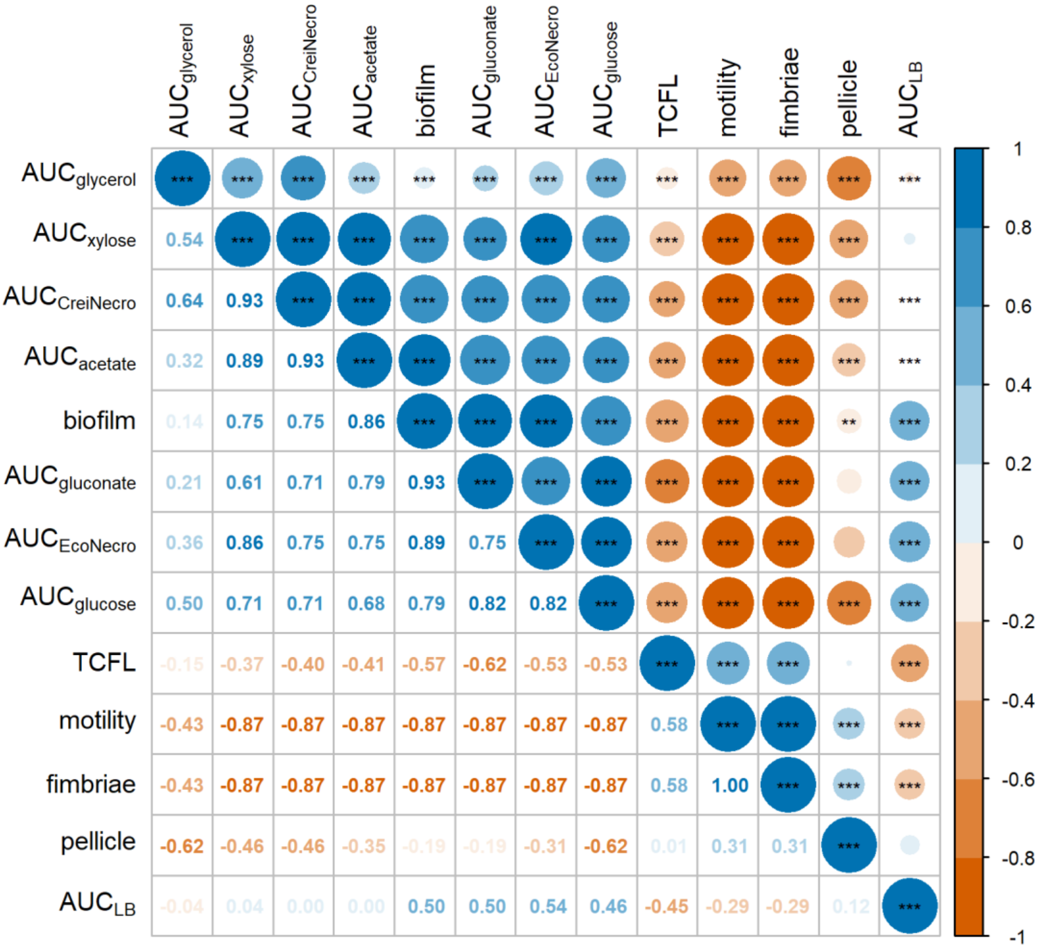


**Supplementary Figure 10.** Heatmap of the correlation matrix between the time to chlorophyll fluorescence loss (TCFL) of co-cultured microbial biospheres and the phenotypes and carbon source utilization capacities of their constituent *Escherichia coli* (*E. coli*) populations. Spearman's rank correlation coefficients (ρ) are displayed in the lower triangle; the corresponding significance levels (indicated by asterisks) are shown in the upper triangle. CreiNecro and EcoNecro denote necromass derived from *Chlamydomonas reinhardtii* and *E. coli*, respectively. AUC (area under the curve) is used as a metric for carbon source utilization capacity. TCFL represents the persistence of the co-culture system. Motility and curli fimbriae expression were assessed qualitatively using motility test media and Congo Red indicator (CRI) agar, respectively, with results on a binary scale[1,2]. For the correlation analysis, biofilm formation capacity was quantified using raw OD570 values from the tissue culture plate (TCP) assay (72-hour data), rather than categorical classifications.[2] Pellicle formation was similarly evaluated qualitatively, following the TCP protocol but with cultures grown in test tubes.[3]

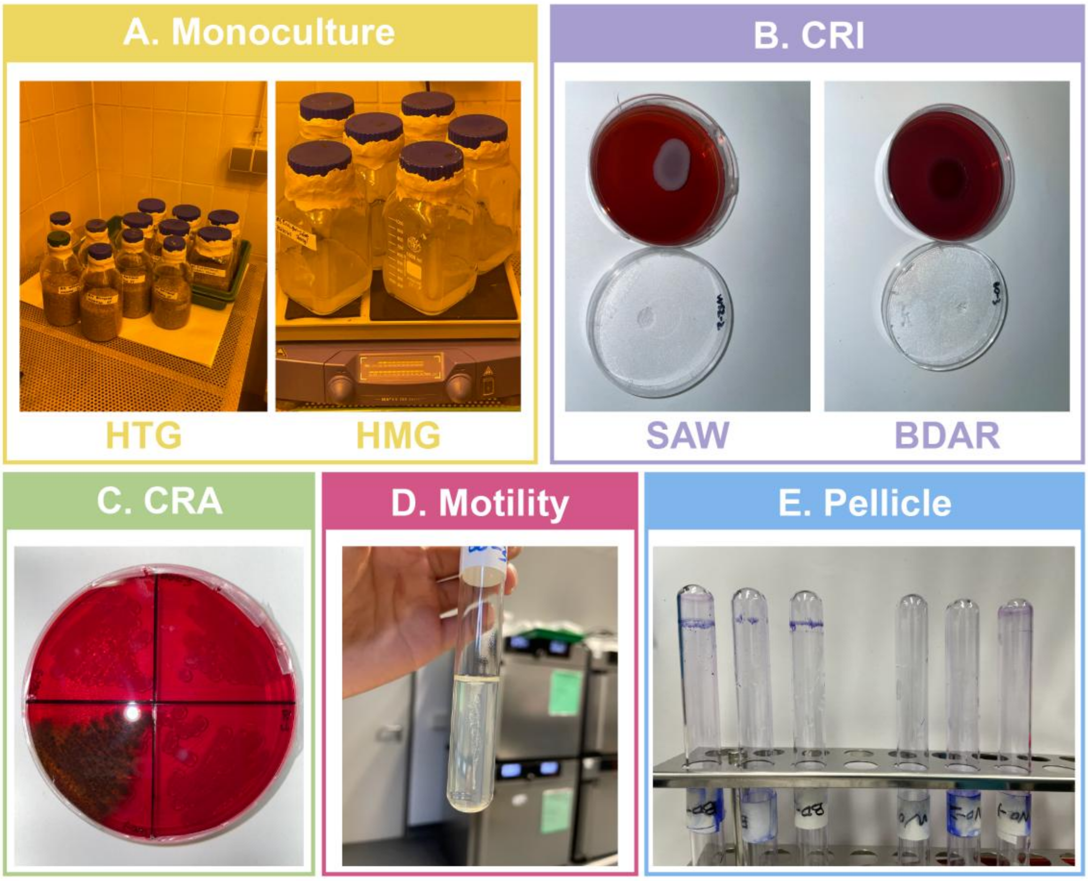


**Supplementary Figure 11.** Photographic displays of (A) monoculture, (B) the CRI assay, (C) the CRA assay, (D) the motility test, and (E) pellicle formation.

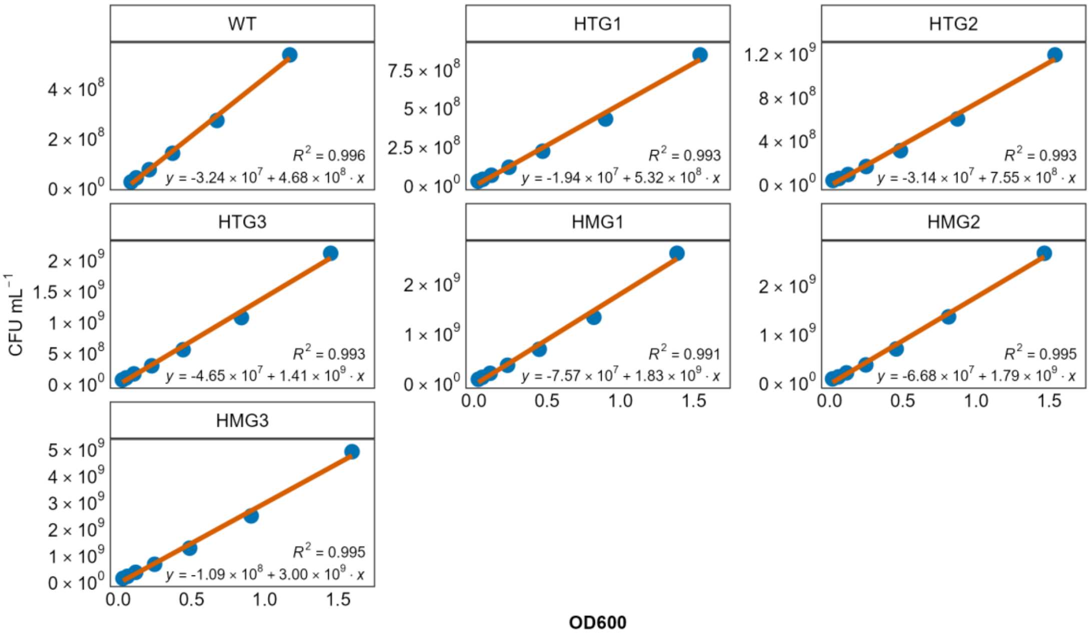


**Supplementary Figure 12.** Standard curves relating CFU $mL^{-1}$ to OD600. The curves were fitted and parameters were derived using ordinary least squares (OLS) regression. The conversion formula for each is indicated in the bottom right corner of the respective panel.

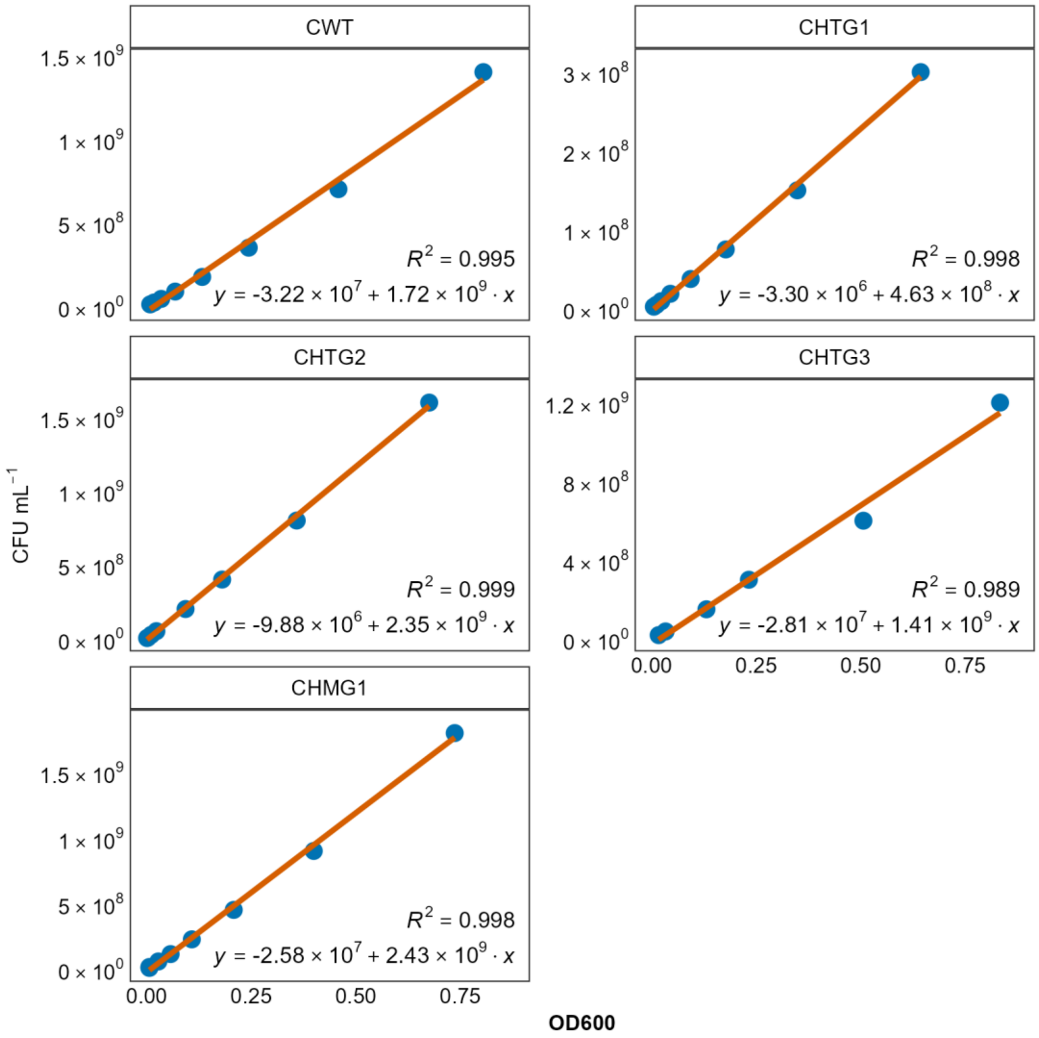


**Supplementary Figure 13.** Supplementary standard curves relating CFU $mL^{-1}$ to OD600. Curves were derived from the five *Escherichia coli* populations successfully recovered at the fourth destructive harvest; the prefix "C" denotes co-culture. Curves were fitted and parameters were derived using ordinary least squares (OLS) regression. The conversion formula for each is indicated in the bottom right corner of the respective panel.

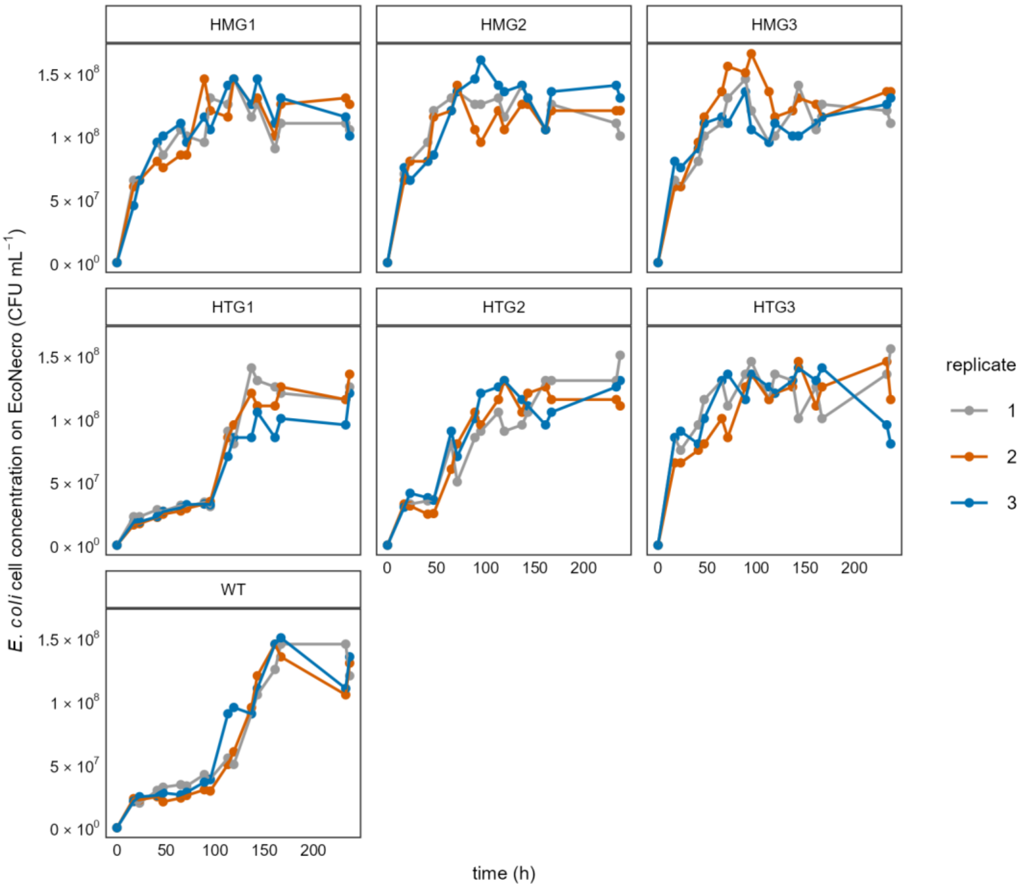


**Supplementary Figure 14.** Growth curves of *Escherichia coli* (*E. coli*) MC4100 populations in M9 minimal medium supplemented with EcoNecro (*E. coli* necromass). Due to the insolubility of necromass particles, the curve was constructed via manual sampling. For each *E. coli* population, three technical replicates were established. Over the 10-day experiment, manual sampling was performed at an average interval of approximately 15 hours, resulting in 17 time points. *E. coli* cell numbers were enumerated using the SP-SDS method.[4]

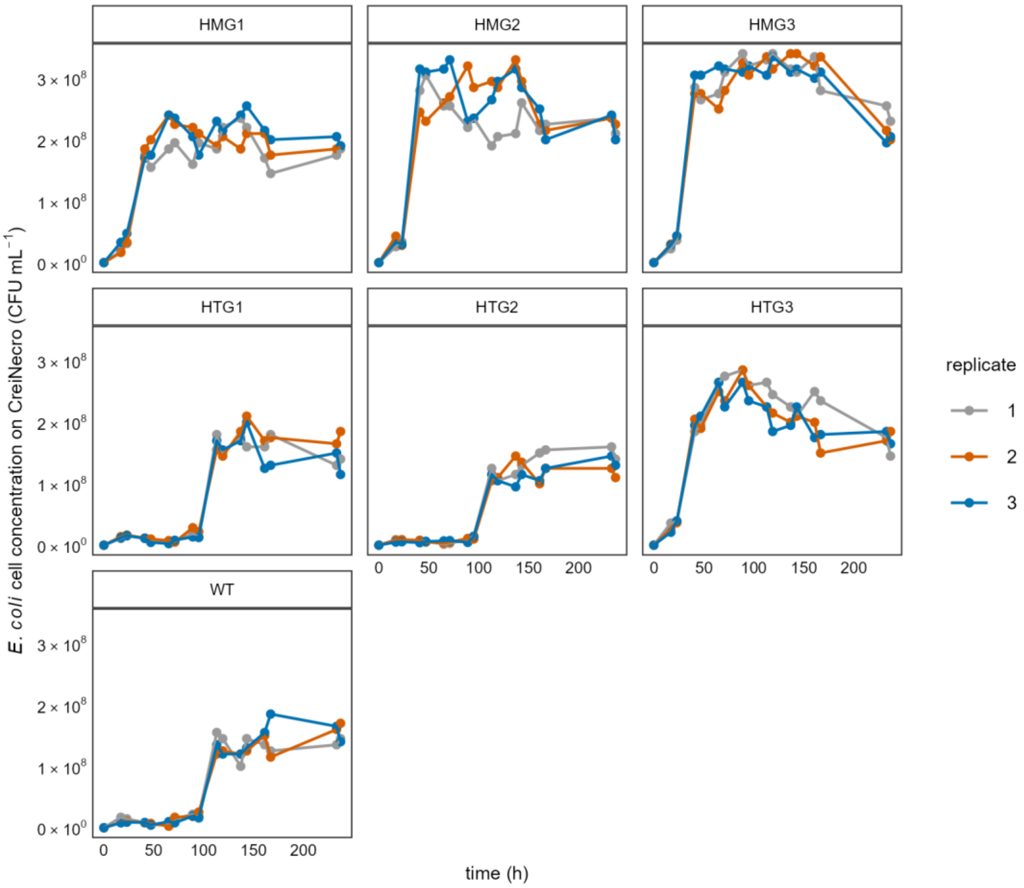


**Supplementary Figure 15.** Growth curves of *Escherichia coli* (*E. coli*) MC4100 populations in M9 minimal medium supplemented with CreiNecro (*Chlamydomonas reinhardtii* necromass). Due to the insolubility of necromass particles, the curve was constructed via manual sampling. For each *E. coli* population, three technical replicates were established. Over the 10-day experiment, manual sampling was performed at an average interval of approximately 15 hours, resulting in 17 time points. *E. coli* cell numbers were enumerated using the SP-SDS method.[4]
